# Photochromic fluorophores enable imaging of lowly-expressed proteins in the autofluorescent fungus *Candida albicans*

**DOI:** 10.1101/2021.02.26.433138

**Authors:** Wouter Van Genechten, Liesbeth Demuyser, Sam Duwé, Wim Vandenberg, Patrick Van Dijck, Peter Dedecker

**Affiliations:** Laboratory of Molecular Cell Biology, Institute of Botany and Microbiology, KU Leuven, 3001 Leuven, Belgium; VIB-KU Leuven Center for Microbiology, KU Leuven, 3001 Leuven, Belgium; Advanced Optical Microscopy Centre, Biomedical Research Institute (BIOMED), Hasselt University, 3500, Hasselt, Belgium; Lab for Nanobiology, Department of Chemistry, KU Leuven, 3001 Leuven, Belgium

## Abstract

Fluorescence microscopy is a standard research tool in many fields, though collecting reliable images can be difficult in systems characterized by low expressions levels and/or high background fluorescence. We present the combination of a photochromic fluorescent protein and stochastic optical fluctuation imaging (SOFI) to deliver suppression of the background fluorescence. This strategy makes it possible to resolve lowly- or endogenously-expressed proteins, as we demonstrate for Gcn5, a histone acetyltransferase required for complete virulence, and Erg11, the target of the azole antifungals agents in the fungal pathogen *C. albicans*. We expect that our method can be readily used for sensitive fluorescence measurements in systems characterized by a high background fluorescence.

**Importance:** Understanding the spatial and temporal organization of proteins-of-interest is key to unravel cellular processes and identify novel possible antifungal targets. Only a few therapeutic targets have been discovered in *Candida albicans* and resistance mechanisms against these therapeutic agents is rapidly acquired. Fluorescence microscopy is a valuable tool to investigate molecular processes and assess the localization of possible antifungal targets. Unfortunately, fluorescence microscopy of *C. albicans* suffers from extensive autofluorescence. In this work we present the use of a photochromic fluorescent protein and stochastic optical fluctuation imaging to enable imaging of lowly-expressed proteins in *C. albicans* through the suppression of autofluorescence. This method can be applied in *C. albicans* research or adapted for other fungal systems allowing the visualization of intricate processes.

## Introduction

Fluorescence imaging has contributed to many discoveries in a wide variety of systems. The microscopic imaging of fungi, for example, has led to a greater understanding of the virulence mechanisms in pathogens such as *C. albicans*. This opportunistic fungal pathogen can cause bloodstream infections, also called candidemia, which has a crude mortality rate of approximately 40% (1). Fluorescence imaging of this organism has been performed on both a macro scale (2) and on a subcellular scale, though both approaches tend to make use of overexpression of specific targets labeled with a range of fluorescent proteins (3-6). Imaging of labeled molecules is often hampered by autofluorescence, which can originate from the presence of many weakly fluorescent molecules such as vitamin B_2_ (11, 12). This vitamin, also termed riboflavin, is synthesized by *C. albicans* upon activation of the PKA pathway (11). Its green-yellow emission is especially problematic when the labeled proteins are expressed at low levels, as is often the case for endogenous expression levels, especially because the brightest and best-maturing fluorescent proteins are typically located in this spectral region. Overexpression of fusion constructs is one way of improving the obtained signal, but may interfere with the cellular phenotype (13) and can lead to accumulation in the endoplasmic reticulum (ER), vacuole and terminal membranes (13).

These limitations pose challenges for many cellular targets. One example is Erg11, a cytochrome P450 (CYP) (16) and target of the first-line antifungal fluconazole (17). It is known that overexpression of CYPs can lead to ER stress (18), and visualization of Erg11 under endogenous expression is therefore important to correctly assess its localization. Previous experiments by our group showed a localization of Erg11 to the ER and plasma membrane in *Saccharomyces cerevisiae* under conditions of overexpression (15). Another example is Gcn5, a histone acetyltransferase involved in hyphae formation and cell wall stress response. Overexpression showed localization to the nucleus during the stationary phase and in the cytosol during the exponential phase (5), though imaging of Gcn5 under endogenous expression did not result in a usable signal.

Recently, we codon-optimized fast-folding Dronpa (ffDronpa), a novel type of ’smart’ fluorescent protein, resulting in the CeffDronpa label for applications in *C. albicans* (4). The fluorescence emission of ffDronpa can be switched ‘on’ and ‘off’ using 405-and 488-nm light respectively (19), enabling applications in diffraction-unlimited fluorescence imaging (20) using techniques such as photochromic stochastic optical fluctuation imaging (pcSOFI), photo-activated localization microscopy (PALM), or Reversible Saturable Optical Fluorescence Transitions (RESOLFT) (21-23). Photochromic SOFI relies on the acquisition of multiple fluorescence images (hundreds or more) from fluorophores that show fluorescence dynamics or ’blinking’. Statistical analysis of the acquired images then leads to a super-resolved image with a resolution enhancement of up to two-or three-fold or more (21, 24).

Since it relies on the analysis of single-molecule fluorescence fluctuations, pcSOFI is highly sensitive to the emission from bright emitters with pronounced intensity fluctuations, such as photochromic fluorescent proteins, but effectively insensitive to emission originating from a large number of weakly autofluorescent molecules, as is typically the case for background emitters.

Other reports have investigated the use of photochromic proteins to reduce background levels, though these approaches often require customization of the instrument so that it can generate specific illumination sequences with precise timing control (28-30). pcSOFI, in contrast, derives its enhancement from the spontaneous blinking of the fluorophores, which means that no adaptation or customization of the instrument is required.

In this contribution, we applied pcSOFI microscopy to samples labeled with CeffDronpa. We find that this strategy results in efficient background rejection and a more straightforward assessment of protein localization without interference of autofluorescence. We also demonstrate the imaging of lowly expressed targets, thereby increasing the set of proteins that can be visualized via labeling of the endogenous population. In particular, we show that Erg11 localizes to the ER and the plasma membrane in *C. albicans*, while a proportion of Gcn5 localizes to the nucleus and the cytosol depending on the growth phase. We expect that our approach can deliver enhanced imaging in a range of organisms.

## Results and Discussion

### Background fluorescence in C. albicans

We first sought to quantify the background fluorescence in *C. albicans* wild type SC5314 relative to *S. cerevisiae* laboratory strain S288C. Flow cytometry measurements show a significant difference (p = 0.003) in the emission of the unlabeled cells, even though both strains have a similar physical size as measured using the forward scatter component. As can be seen from figure 1A, the background signal of *C. albicans* is almost twofold higher, indicating that fluorescence imaging of *C. albicans* in a similar spectral window is more challenging. This bright autofluorescence is omnipresent in filamentous fungi where lipofuscin, ergosterol, riboflavin, flavin adenine dinucleotide, and cell wall components significantly contribute to the background signal (11, 25-27, 31).

**Figure 1.**
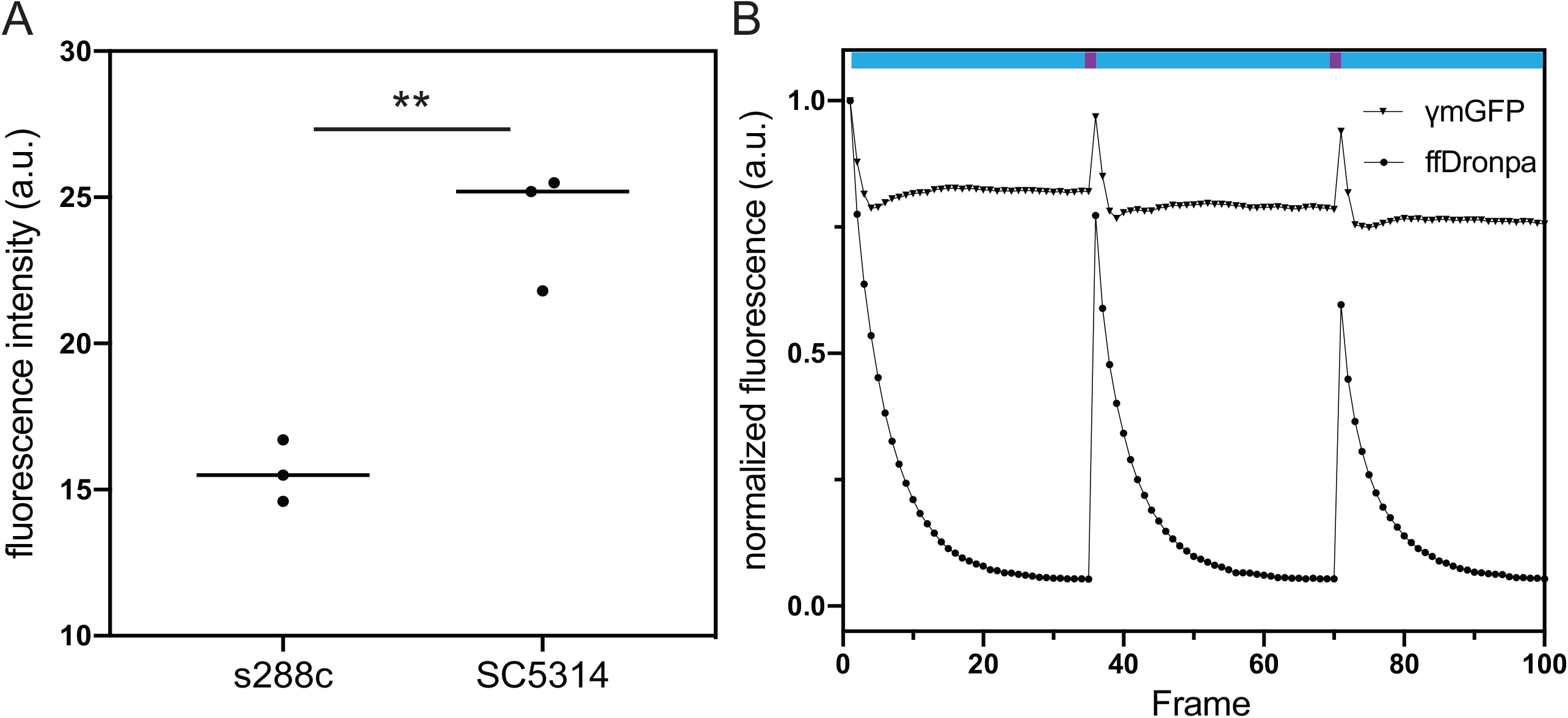
A) Autofluorescence of *S. cerevisiae* diploid strain S288C and *C. albicans* wild type SC5314 measured using flow cytometry. Cells were grown overnight on LoFlo medium to the exponential phase (4-5 hrs) before measurement. Asterisks denote significance level with ** corresponding to p < 0.01 as calculated using Student’s t-test. B) *In vivo* photoswitching assessment of *C. albicans* overexpressing CeffDronpa or ymGFP. Fluorescent proteins were cycled by exposing the cells to one pulse of 405-nm laser light (purple bar) to induce switching to the fluorescent state before capturing 35 images using 488-nm excitation (cyan bar).

### In vivo reversible switching of Candida-enhanced ffDronpa

We then verified whether CeffDronpa retains its photochromism in *C. albicans*. We overexpressed the protein and found that it is able to switch reversibly between the dark and the bright state as indicated in figure 1B. The initial fluorescence of the cells is relatively high and drops quickly to a plateau approximately equal to background fluorescence upon applying 488 nm illumination, resulting in a large contrast between the on-and off-state of the FP *in vivo*. A subsequent 405 nm pulse is able to recover the fluorescence to approximately 75% of the maximum fluorescence of the previous cycle, suggesting some photodestruction of the label. Thus, *in vivo* CeffDronpa (pKa 5.0) displays robust photochromism even though the cytosolic pH of *Candida albicans* ranges from 6 to 7, which is more acidic than the mammalian cytosolic pH of 7.1-7.2 (32). We also included γmGFP-expressing cells as a reference. We find that the resulting cells show a ±25% higher fluorescence brightness, but do not show the pronounced modulation pattern displayed by cells expressing CeffDronpa.

### Localization of Gcn5 to the nucleus and the cytosol

Gcn5 is a catalytic subunit of the histone acetyltransferase complex SAGA, which is involved in one of the main virulence factors of *C. albicans*. Deletion of *GCN5* rendered *C. albicans* non-virulent in a mouse tail vein infection model (5). Imaging of a Gcn5–GFP fusion construct under constitutive expression from the *ADH1* promoter (overexpression) showed a nuclear localization in the stationary growth phase and a cytoplasmatic localization in the exponential growth phase (5). However, no signal was observed from expression of the fusion construct under the native promotor, rendering the localization of Gcn5 at endogenous levels uncertain (5).

We constructed a Gcn5-CeffDronpa fusion protein under control of the endogenous promotor and visualized its expression in live cells. Figure 2 shows the averaged (non-pcSOFI) signal from the green channel as well as an overlay with the Nucblue™ nuclear marker. This averaged image was obtained by combining the first 15 images in the SOFI image stack, and can be considered to be a classical wide-field image of the same region of the same sample. The non-SOFI images (panels A, E) show some brighter features and a high degree of autofluorescence, comparable to images acquired on cells that did not express any fluorescent proteins (panels I, M). We concluded that it is difficult to distinguish relevant features from the cellular autofluorescence, confirming the difficulty of observing Gcn5 at endogenous levels.

**Figure 2.**
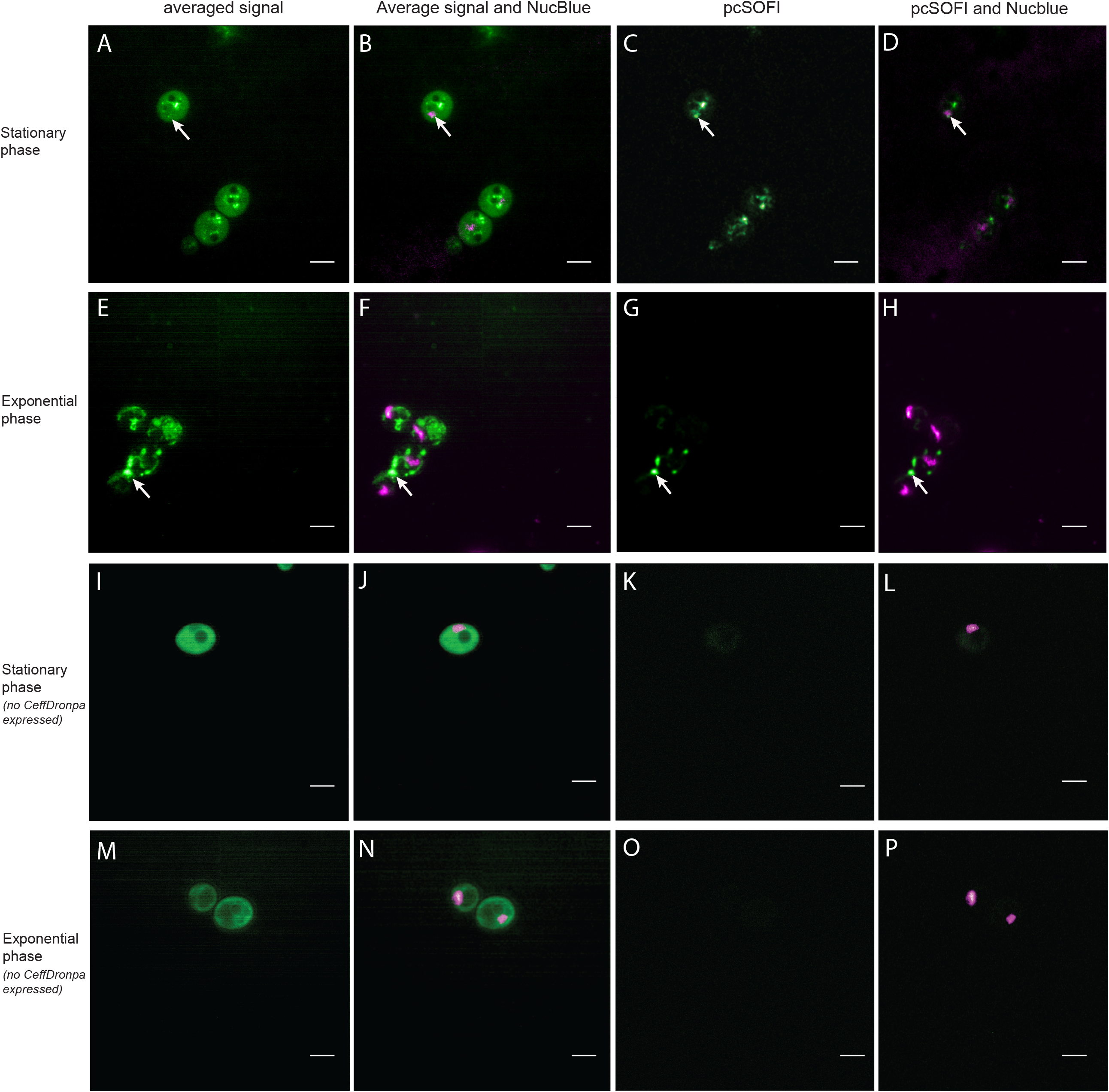
Imaging of Gcn5 tagged with CeffDronpa in the stationary phase (A-D) and the exponential phase (E-H) using pcSOFI. White arrows indicate points-of-interest in both the averaged widefield and the pcSOFI processed image. A,E) the averaged image of a stack obtained in the green channel by exciting cells using 488-nm light. B,F) Overlay of the averaged image with the Nucblue staining in magenta. C,G) pcSOFI postprocessing of the same stack. D,E) Overlay of the pcSOFI resulting image with the NucBlue staining in magenta. Panels I-P contain the Imaging of the negative control samples for the Gcn5 – CeffDronpa measurements in the stationary phase (I-L) and the exponential phase (M-P). I,M) the averaged image of a stack obtained in the green channel by exciting cells using 488-nm light. J,N) Overlay of the averaged image with the Nucblue staining in magenta. K,O) pcSOFI postprocessing of the same stack. L,P) Overlay of the pcSOFI resulting image with the NucBlue staining in magenta. Scalebars indicate 5 μm.

We then applied our pcSOFI analysis to these samples, leading to the images shown in panels C, D, G and H. Compared to the conventional imaging, the pcSOFI images show more cellular structuring and features and do not show the unstructured background emission. Some of these features could be partially observed already in the conventional imaging, though the pcSOFI imaging improved the contrast (white arrows). A representative dataset is shown for both the stationary and exponential phase. Additional datasets of the biological replicates in each growth condition are available (10.5281/zenodo.4265989).

During the stationary phase a proportion of the signal obtained from the Gcn5 fusion appears to co-localize to the nucleus (panel D), while exponentially growing cells show distinct structures without indications of overlap with the nuclear staining (panel G and H). Negative control images of cells stained with NucBlue™ but not expressing CeffDronpa show autofluorescence in the average images, but no pcSOFI signal in both the stationary and the exponential phase (panels K, O). This shows that neither the NucBlue™ staining nor the cellular autofluorescence lead to the fluorescence dynamics detected by the pcSOFI signal.

### Localization of Erg11 to the ER and plasma membrane

The most widely used class of antifungals, the azoles, target the biosynthesis of ergosterol. More specifically, azoles target the lanosterol-14-alfa-demethylase enzyme encoded by *ERG11*. This key enzyme in the ergosterol biosynthesis pathway is therefore a frequently studied protein which can acquire multiple point mutations that confer resistance to azoles. In *S. cerevisiae* it was shown to localize both to the endoplasmic reticulum and the plasma membrane via Focused Ion Beam Scanning Electron Microscopy (FIB SEM) (15). The extensive sequence similarity between *Sc*Erg11 and *Ca*Erg11 suggests a similar localization may occur in *C. albicans* (33).

Figure 3 shows images obtained on a fusion construct between Erg11 and CeffDronpa expressed at endogenous levels, confirming the results from *S. cerevisiae* and the prediction for *C. albicans*. The averaged signal images (panel A, C) show no visible structuring on account of the autofluorescence which drowns out the CeffDronpa signal. In the pcSOFI processed images (panel B, D), a clear localization to the ER, in the form of a distinct ring around the nucleus, and the plasma membrane is observed. The administration of fluconazole, which was shown to increase the autofluorescence background of SC5314 (supplementary figure 1), did not appear to influence the pcSOFI imaging. No pcSOFI signal was observed in control strains not expressing CeffDronpa (Panel F). Data available at Zenodo 10.5281/zenodo.4265989.

**Figure 3.**
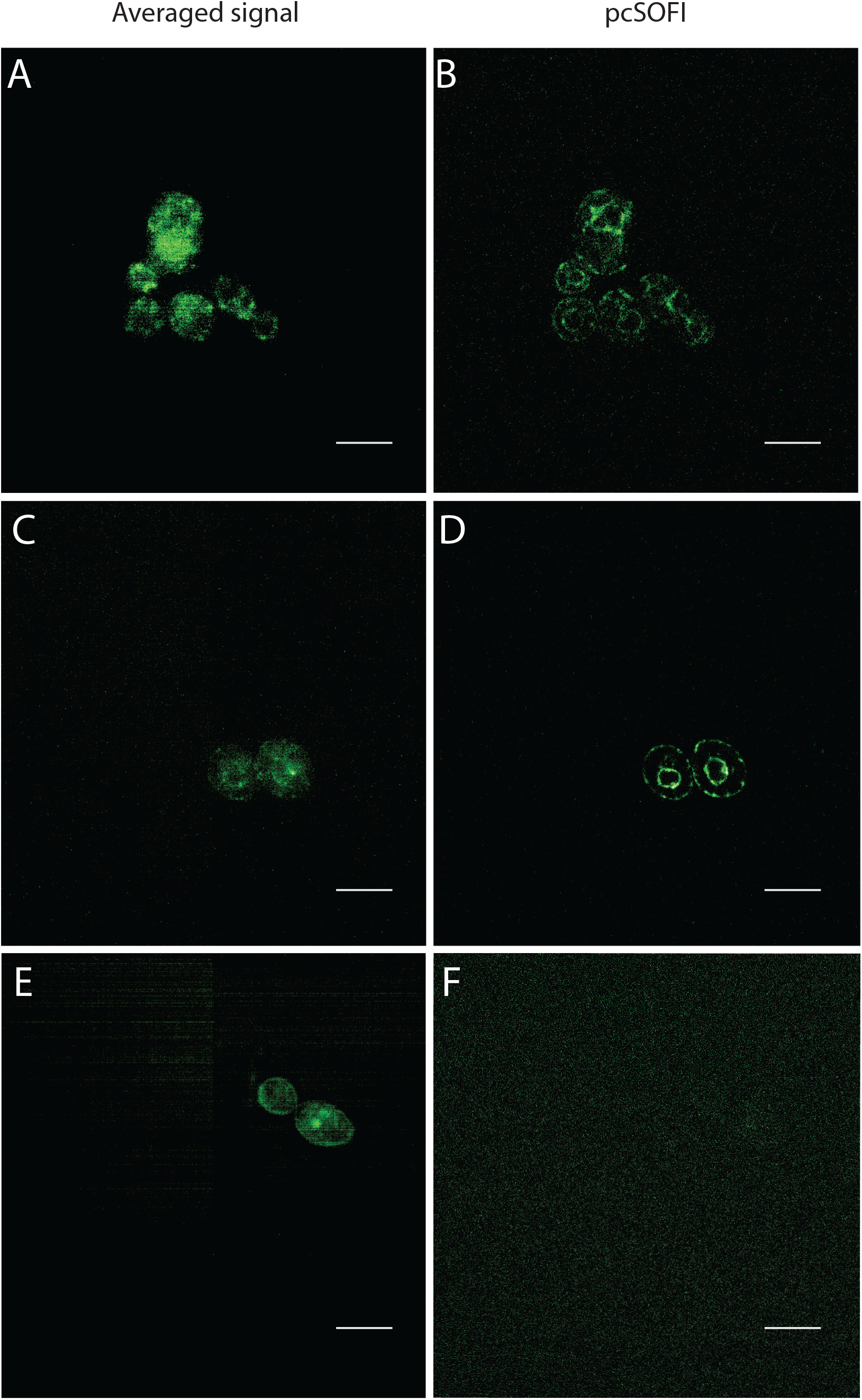
Imaging of Erg11 tagged with CeffDronpa. Cells were grown overnight on LoFlo medium to the exponential phase before measurement. Averaged images of Erg11 tagged with CeffDronpa to provide an indication of a widefield image are depicted in panel A and C. pcSOFI postprocessed images are depicted in panel B and D. The averaged image of a negative control sample is depicted in panel E. The pcSOFI postprocessed image of the negative control sample is depicted in panel F. The slight striping visible in panel A, C and E is due to patterning of the sCMOS camera which becomes visible due to the low light levels, and is not a feature of the sample. Scalebars indicate 5 μm.

These two examples show that the removal of the background autofluorescence and thus the increase in SNR allow for visualization of low abundant targets in living cells. A disadvantage of the imaging is that it requires higher light doses associated with single-molecule detection, and that the temporal resolution is decreased since multiple images must be acquired. We required 18 s to acquire a single pcSOFI image at the expression levels observed here, though this could be accelerated by either lowering the number of collected frames and/or a employing a shorter exposure time at the cost of the resulting SNR.

In conclusion, our work shows that the combination of photochromic fluorophores with pcSOFI allows the reduction of non-label emission in fluorescence microscopy without the customization of the microscope. We demonstrated the potential of this approach by generating endogenous fusion constructs with CeffDronpa and visualizing the resulting fluorescence distribution in *C. albicans*. We propose that this method is a valuable strategy for applications in other systems showing high autofluorescence levels.

## Materials and methods

### Strain construction

For the endogenous tagging of Gcn5 and Erg11, we constructed a pFA6-based plasmid containing the codon-optimized ffDronpa, CeffDronpa. From the CeffDronpa gBlock (IDT) we amplified a Gibson insert using primers listed in supplementary table 1. The pFA6 plasmid, containing a nourseothricin marker, was digested with *Pst*I and *Sma*I before insertion of CeffDronpa. From this pFA6-CeffDronpa plasmid, we created linear PCR-fragments with 100 bp homologous overhang for recombination with the 3’-end of the *GCN5* or *ERG11* gene, without the stop codon, and the 5’-start of its terminator. Between the *GCN5* or *ERG11* gene and the start of the FP, we included a linker sequence containing a triple repetition of glycine and alanine (GAGAGA). These linear fragments were transformed into the wild type *C. albicans* strain, SC5314. Genomic DNA was sequenced to confirm correct integration of CeffDronpa.

### Photoswitching assessment

*Candida* enhanced ffDronpa and γmGFP expressing strains that have been used in the photoswitching assessment were published previously (4) and grown overnight on Low Fluorescent medium (LoFlo) containing 0.69%(w/v) Yeast Nitrogen Base (without amino acids, folic acid and riboflavin), 0.25%(w/v) ammonium sulphate, 0.079%(w/v) complete supplement mixture (CSM; MP Biomedicals) and 2%(w/v) glucose. The strains were brought to an optical density at 600 nm (OD_600_) of 0.2 in fresh Low Fluorescent medium and grown to the exponential phase (4-5 hrs) before measurement of photoswitching. The microscopic setup used for photoswitching and the following imaging experiments, is a Nikon Ti2 epifluorescence microscope equipped with a Nikon 100x CFI apo objective (NA = 1.49). Lasers used for the photoswitching imaging are 405 and 488nm Oxxius lasers. To assess the photoswitching capacity of *in vivo* CeffDronpa, a cycle of on- and off-switching was induced consisting of a 405-nm pulse, followed by acquisition of 35 images using 488-nm excitation and detection with a 540/30 bandpass filter. Fluorescence images were collected with a Hamamatsu imageEM X2. The same measurement regime was applied to γmGFP expressing cells and wild type SC5314. The fluorescence intensity of 10 cells for each strain was measured by manually selecting regions-of-interest (ROI) and normalized to the first frame.

### Flow Cytometry measurement of background signal

The background signal of three replicates of the *C. albicans* wild type strain SC5314 and a diploid *Saccharomyces cerevisiae* strain, S288C, were measured using Flow Cytometry. Strains were grown overnight in YPD before bringing them to an optical density of 0.2. Cultures were grown for approximately 5 hours in LoFlo medium in a 30°C shaking incubator before measuring 100 000 cells with a BD influx flow cytometer with 488-nm excitation and a bandpass emission filter 530/40. Mean fluorescence intensity was calculated for each strain using Flowjo^™^ (34) and a Student’s t-test was performed using GraphPad Prism.

### pcSOFI imaging

#### Gcn5 imaging and pcSOFI processing

The experiment was carried out in a way to resemble the first publication reporting the localization of Gcn5 (5). Cells were grown overnight in LoFlo medium until stationary phase. Part of this culture is diluted to an optical density of 0.2 in fresh LoFlo medium. These cells were grown for 4-5 hours at 30°C until mid-exponential phase before staining the nucleus with NucBlue™.

We first imaged the NucBlue™ staining using excitation with 405-nm light and a 480/40 emission filter, subsequently we acquired at least 1500 frames of CeffDronpa tagged to Gcn5 using 488-nm excitation. Emission was recorded through a bandpass filter 540/30. To construct the SOFI image using the Localizer package in IgorPro (35), 600 frames where used, dropping the initial 15 frames to exclude non-stationary signal. The images produced where checked for artifacts using SOFIevaluator (36). Since the SNR of the images was limited, the image was subsequently convolved with a gaussian (FWHM 112,5nm) to further smooth out the background noise and bring out the labelled features. Average images of frame 16 to 30 were also calculated to serve as an indication of the CeffDronpa abundance and background intensities.

#### Erg11 imaging and pcSOFI processing

Cells were grown overnight in LoFlo medium until stationary phase before bringing this culture to an OD_600_ of 0.2. To simulate experimental conditions under which research is performed within the susceptible-dose dependent range of concentrations, we administered 16 μg/mL fluconazole to the culture (37, 38). Cells were grown at 30°C for 5 hours before visualization using the same microscope setup as described earlier. The pcSOFI image analysis was performed using the Localizer package in IgorPro (35). The first 5 frames were not utilized in the construction due to the initial non-stationary behavior. Frames 6 till 1000 were utilized for the reconstruction. Average images of the first 15 frames were also calculated to serve as an indication of the CeffDronpa abundance and background intensities.

## Acknowledgement

This work was supported by the Research Foundation Flanders (FWO Vlaanderen) [1S01817N to W.V.G., G062616N to P.V.D. and P.D.] and ERC Starting Grant 714688 *NanoCellActivity*. We thank Nico Vangoethem for the generous help in the creation of the figures.

## Supplementary information

Fig. S1. Autofluorescence measurement using flow cytometry of 100 000 cells of SC5314 grown in LoFLo medium with and without 16μg/mL fluconazole to exponential phase. Student’s t-test performed with Graphpad Prism gave a significant difference in autofluorescence between these growth conditions (p=0.0237) indicated with an asterisk.

Table S1. Primers for the construction of the pFA6-CeffDronpa plasmid and endogenous tags using the PCR-mediated integration of the fluorescent protein.

